# Predicting symptom severity in autism spectrum disorder based on cortical thickness measures in agglomerative data

**DOI:** 10.1101/039180

**Authors:** Elaheh Moradi, Budhachandra Khundrakpam, John D. Lewis, Alan C. Evans, Jussi Tohka

## Abstract

Machine learning approaches have been widely used for the identification of neuropathology from neuroimaging data. However, these approaches require large samples and suffer from the challenges associated with multi-site, multi-protocol data. We propose a novel approach to address these challenges, and demonstrate its usefulness with the Autism Brain Imaging Data Exchange (ABIDE) database. We predict symptom severity based on cortical thickness measurements from 156 individuals with autism spectrum disorder (ASD) from four different sites. The proposed approach consists of two main stages: a domain adaptation stage using partial least squares regression to maximize the consistency of imaging data across sites; and a learning stage combining support vector regression for regional prediction of severity with elastic-net penalized linear regression for integrating regional predictions into a whole-brain severity prediction. The proposed method performed markedly better than simpler alternatives, better with multi-site than single-site data, and resulted in a considerably higher cross-validated correlation score than has previously been reported in the literature for multi-site data. This demonstration of the utility of the proposed approach for detecting structural brain abnormalities in ASD from the multi-site, multi-protocol ABIDE dataset indicates the potential of designing machine learning methods to meet the challenges of agglomerative data.

## Introduction

Autism Spectrum Disorder (ASD) is a developmental disorder characterized by impairments in social interaction and communication, restricted interests, and repetitive patterns of behaviour (Lord and Jones, 2012; Wing, 1997; Gillberg, 1993). The definition admits substantial behavioural heterogeneity (Georgiades et al., 2013); ASD is, in fact, a family of developmental disorders with unique, but related, phenotypes, with a variety of genetic associations (Devlin and Scherer, 2012). Moreover, ASDs are developmental disorders, and the behavioural abnormalities evolve over time (Gotham et al., 2012; Szatmari et al., 2015), adding to the apparent heterogeneity. This large behavioural heterogeneity appears to be paralleled by a wide array of neuroanatomical abnormalities, which also evolve over development (Zielinski et al., 2014;Wolff et al., 2014). Almost every brain region has been implicated in autism, including subcortical (Jacobson et al., 1988; Cerliani et al., 2015) and cerebellar regions (Bauman, 1991; Fatemi et al., 2002), gray-matter and white-matter (Barnea-Goraly et al., 2004; Rojas et al., 2006), and regions of all lobes of the cerebrum (Zilbovicius et al., 2000; Courchesne et al., 2011; Lewis et al., 2013, 2014). Indeed, the neuroanatomical heterogeneity is so great that replication of results across studies is rare. The inconsistencies in findings are likely primarily due to the small sample sizes used in most studies, in combination with the large behavioural heterogeneity, as well as measurement related differences (Auzias et al., 2014, 2016; Castrillon et al., 2014). Thus, there is an urgent need for larger sample sizes, if we are to discover clinically useful information (Amaral et al., 2008; Auzias et al., 2014, 2016; Lefebvre et al., 2015). Large samples may allow the extraction of core neuroanatomical abnormalities from the noise introduced by the heterogeneity of the disorder. Such abnormalities could serve as biomarkers, and could provide insight into the causes of the disorder, and potential interventions.

However, datasets collected by a single site are not sufficient in size to achieve such goals (albeit making exact claims about the required dataset size is a complex matter and depends on the goals of study (Button et al., 2013)). Further, there are limited publicly available data from multi-site studies utilizing a single scanner type with the same acquisition protocol across sites. But, so-called ‘big data’ has come to neuroscience, including for the study of ASD. There are currently multiple initiatives to bring together neuroimaging data from multiple sites, acquired on multiple types of scanners, and with differing protocols. The Autism Brain Imaging Data Exchange (ABIDE)^1^ is one such initiative (Di Martino et al., 2014). ABIDE provides previously collected datasets composed of both MRI data and phenotypic information from 16 different international sites for over 1100 individuals, approximately half of whom are typically developing (TD) and half have been diagnosed with ASD. This sample size, which is more than an order of magnitude larger than that used in most single-site studies, provides the power needed to identify neuroanatomical abnormalities related to ASD. But, the multisite, multi-protocol aspect of the data introduces additional heterogeneity. Indeed, previous studies using the ABIDE data have shown that acquisition site has significant effects on basic image properties (Nielsen et al., 2013; Castrillon et al., 2014). This further exacerbates the problem of identification of core neuroanatomical abnormalities in this extremely heterogeneous data. The between-site heterogeneity constitutes the main technical challenge in the current work (Auzias et al., 2014), and the solution that we offer is a contribution applicable not only to the ABIDE dataset, but to any neuroimaging data agglomeration.

The solution to the problem lies in finding a new common space within different datasets for reduction of between-site variation. Techniques for achieving this are often referred to as *domain adaptation* (Jiang, 2008; Pan and Yang, 2010). Domain adaptation is a new branch of machine learning techniques that seeks to improve the similarity of the data from different sources with mismatched distributions. We utilize these domain adaptation machine learning algorithms to address the problem that arises in the situation where the data distribution changes across different acquisition sites. We apply this approach to the ABIDE data to identify neuroanatomical abnormalities associated with symptom severity in ASD. Between-sites variance in neuroimaging studies is commonly handled by regressing out the site identity from the imaging data in a voxel-wise manner before performing analysis (Gupta et al., 2015) and similar methods have been adapted for machine learning analysis with limited success (Kostro et al., 2014). Instead, here we propose a novel approach for reducing between-sites variability by projecting data from different sites into a new, common space in a way that effectively reduces nuisance variation between the data from different sites. The current approach for dealing with the site effect is novel in the context of multi-site imaging studies, and for the estimation of severity scores in ASD patients.

The great majority of ASD studies have focused on identifying group differences between typically developing individuals and those with ASD, or conversely, training classifiers to distinguish between these groups (Ecker et al., 2010; Nielsen et al., 2013; Wang et al., 2015). But, perhaps the largest source of heterogeneity is associated with the severity of the disorder. In fact, both individuals with ASD as well as those deemed to be typically developing display a wide range of symptoms of autism in a variety of behaviours. This variability may mask neural abnormalities associated with these symptoms, and limit the success of attempts to classify an individual based on their neuroimaging data. Approaches which relate dimensional measures of symptoms to measures of neuroanatomy appear more useful than those which aim only to identify abnormalities associated with a diagnosis of ASD (Sato et al., 2013; Schumann et al., 2009). Thus, in this work we take this latter approach. We design a model to estimate symptom severity scores derived from the Autism Diagnostic Observation Schedule (ADOS) from cortical thickness measurements. We are motivated by evidence that local cortical thickness measures provide an index of the maturation of cortex and cortico-cortical connectivity (Shaw et al., 2008; Raznahan et al., 2011), and that ASD may be characterized by delayed maturation (Webb et al., 2011; Johnson et al., 2015).

Our proposed method for estimation of the severity score consists of two main stages: a domain adaptation stage that uses partial least squares regression (PLS) with sites as response variable, and the learning stage which consists of the combination of two different regression methods, i.e. support vector regression (SVR) and elastic-net penalized linear regression (LR). We evaluate the reliability of the model across a multisite dataset without standardization of the acquisition protocol across sites, and the effect of each part of the algorithm.

## Materials and methods

### 2.1. ABIDE data

The data used in this study were from the ABIDE dataset (Di Martino et al., 2014). ABIDE is a publicly available dataset that involved 16 international sites, from 532 individuals with ASD and 573 typical controls, yielding 1112 datasets composed of MRI (functional and structural) and phenotypic information for each subject. The sequence parameters as well as type of scanner varied across sites, though all data were collected with 3 Tesla scanners. The scan procedures and parameters are described on the ABIDE website ^2^.

### 2.2. Image preprocessing

The T1-weighted volumes were processed with CIVET, a fully automated structural image analysis pipeline developed at the Montreal Neurological Institute. CIVET corrects intensity non-uniformities using N3 (Sled et al., 1998); aligns the input volumes to the Talairach-like ICBM-152-nl template (Collins et al., 1994); classifies the image into white matter, gray matter, cerebrospinal fluid, and background (Zijdenbos et al., 2002; Tohka et al., 2004); extracts the white-matter and pial surfaces (Kim et al., 2005); and warps these to a common surface template (Lyttelton et al., 2007). Cortical thickness (CT) is measured in native space using the linked distance between the two surfaces at 81,924 vertices. The thickness map was then blurred to impose a normal distribution on the corticometric data, and to increase the signal to noise ratio; a 30-millimeter full width at half maximum surface-based diffusion smoothing kernel was used.

Quality control (QC) of the CIVET results was performed by two independent reviewers. Data with artifacts due to motion, low signal to noise ratio, hyperintensities from blood vessels, or poor placement of the gray or white matter (GM and WM) surfaces for any reason were excluded. 215 subjects with ASD were excluded in the QC.

### 2.3. Subjects

After image preprocessing and the QC, the number of ASD subjects reduced from 532 to 317 from 16 different sites. Next, we excluded ASD subjects with missing ADOS total and module information and then we included only subjects from sites containing at least 20 subjects. The remaining 156 subjects were from 4 different sites (NYU, PITT, TRINITY, USM) which were used for estimating severity score. Details of the characteristics of the ABIDE samples used in this work are presented in Table 1. The subject IDs of the included subjects can be found in the supplement.

**Table 1.** Subject demographics; The values are site-wise averages and the values in parentheses provide standard deviations.

| Site | NYU | PITT | TRINITY | USM |
| --- | --- | --- | --- | --- |
| No. of subjects | 72 | 20 | 23 | 41 |
| Males/Females | 61/11 | 17/3 | 23/0 | 41/0 |
| Full Scale IQ | 107.14 (16.64)<br>Range: 76-148 | 112 (13.51)<br>Range: 86-131 | 108.83 (15.23)<br>Range: 72-135 | 102 (17.05)<br>Range: 65-132 |
| Verbal IQ | 105.64 (16.53)<br>Range: 73-139 | 109.60 (12.56)<br>Range: 89-132 | 107.96 (14.45)<br>Range: 85-135 | 98.51 (19.20)<br>Range: 55-130 |
| Performance IQ | 107.58 (17.12)<br>Range: 72-149 | 111.05 (13.53)<br>Range: 87-128 | 107.36 (15.33)<br>Range: 63-131 | 105.15 (17.11)<br>Range: 72-133 |
| Age | 14.82 (7.09)<br>Range: 7-39 | 17.65 (5.84)<br>Range: 9-32 | 17.36 (3.63)<br>Range: 12-26 | 24.61 (8.05)<br>Range: 14-50 |
| ADOS total | 11.25 (4.06)<br>Range: 5-22 | 11.75 (2.97)<br>Range: 7-18 | 10.57 (2.94)<br>Range: 7-17 | 13.22 (3.34)<br>Range: 6-21 |
| Severity score | 6.32 (2.13)<br>Range: 2-10 | 6.70 (1.56)<br>Range: 4-9 | 5.70 (1.82)<br>Range: 3-9 | 7.38 (1.56)<br>Range: 3-10 |

### 2.4. Severity score

This work studies the relation between cortical thickness and measures derived from the Autism Diagnostic Observation Schedule (ADOS) (Lord et al., 2000). The ADOS is a semi-structured assessment of communication, social interaction, and stereotypical behaviours for individuals with autism or other pervasive developmental disorders. The ADOS applies to individuals ranging from nonverbal to verbally fluent, and ranging from infants to adults. But different ADOS modules are utilized, depending on the individual’s developmental and language level, and the scores from different modules are not directly comparable. In order to achieve comparability across modules, the ADOS scores must be transformed to calibrated severity scores (Gotham et al., 2009).

The ABIDE data provides the calibrated severity scores for some but not all subjects; and for those without calibrated severity scores, the information necessary to compute calibrated severity scores is also missing. But a proxy calibrated severity score can be derived from the available ADOS measures. A two-step procedure is used to derive the calibrated severity scores: (i) a weighted sum of ADOS item scores is computed, with the weights determined by Gotham et al. (2007); (ii) the calibrated severity score is retrieved from a lookup table provided by Gotham et al. (2009), which is indexed with the individual’s age, the ADOS module used, and the weighted sum from step (i). For those cases in which ABIDE provides both the total of the social and communication ADOS scores and the weighted sum of the ADOS item scores, the difference between the two is small. We thus approximate the calibrated severity scores by substituting the total of the social and communication ADOS scores for the weighted sum of the ADOS item scores in the first step of the procedure. Our proxy of the calibrated severity score is then arrived at by using the lookup table from Gotham et al. (2009) together with the total of the social and communication ADOS scores, the ADOS module used, and the individual’s age. We investigate the relation between cortical thickness and these proxy calibrated severity scores. Note that one reason for transforming the ADOS scores into calibrated severity scores is to remove effects of the subject demographics, such as age, thus making the calibrated severity scores to more truly reflect the disease severity.

This proxy of the calibrated severity score is discussed in greater detail in the supplementary material. There, for comparison, we also report the experiments of cortical thickness based prediction of the total of the social and communication ADOS scores, using the information of which ADOS modules were used. Severity scores of the included subjects can be found in the supplement.

### 2.5. Overview of methodology

The generic structure of the proposed method is illustrated in Figure 1. The method is divided into two main stages: 1) the domain adaptation stage and 2) the learning stage. In the domain adaptation, first, the cortical thickness measures along cortex were divided into separate regional subsets according to the Automated Anatomical Labelling (AAL) atlas. Each regional subset contains only the vertices belonging to one AAL cortical region. In order to reduce the between-sites variability, we performed PLS based domain adaptation for each subset separately (Section 2.6). This resulted in 78 region-specific site-adapted subsets of cortical thickness components (Figure 1 A) with the same, fixed number of components (25) for each region, thus reducing regional cortical thickness measures into 25 features per a region and a subject. The domain adaptation was performed in an unsupervised manner in all subjects before dividing data into training and test sets. Note that we did not use the severity score (label information) or any kind of cognitive information of the subjects in this stage and only the site information was used as the response variable. This is termed unsupervised domain adaptation, but since all the cortical thickness data is used, the whole learning process becomes transductive that is typical for domain adaptation algorithms (Gong et al., 2012). We stress that the label information was not used so this does not lead to double-dipping. For a clear explanation of this fact and the differences between transductive and inductive machine learning algorithms, we refer to Gammerman et al. (1998). It is important to note that the division of the cortical thickness measures into regional subsets must be done before the PLS-based domain adaptation stage as otherwise the PLS components will not be regionally specific. Also, we need a large enough number of subjects from each site to be able to recognize the possible site-differences.

**Figure 1:**
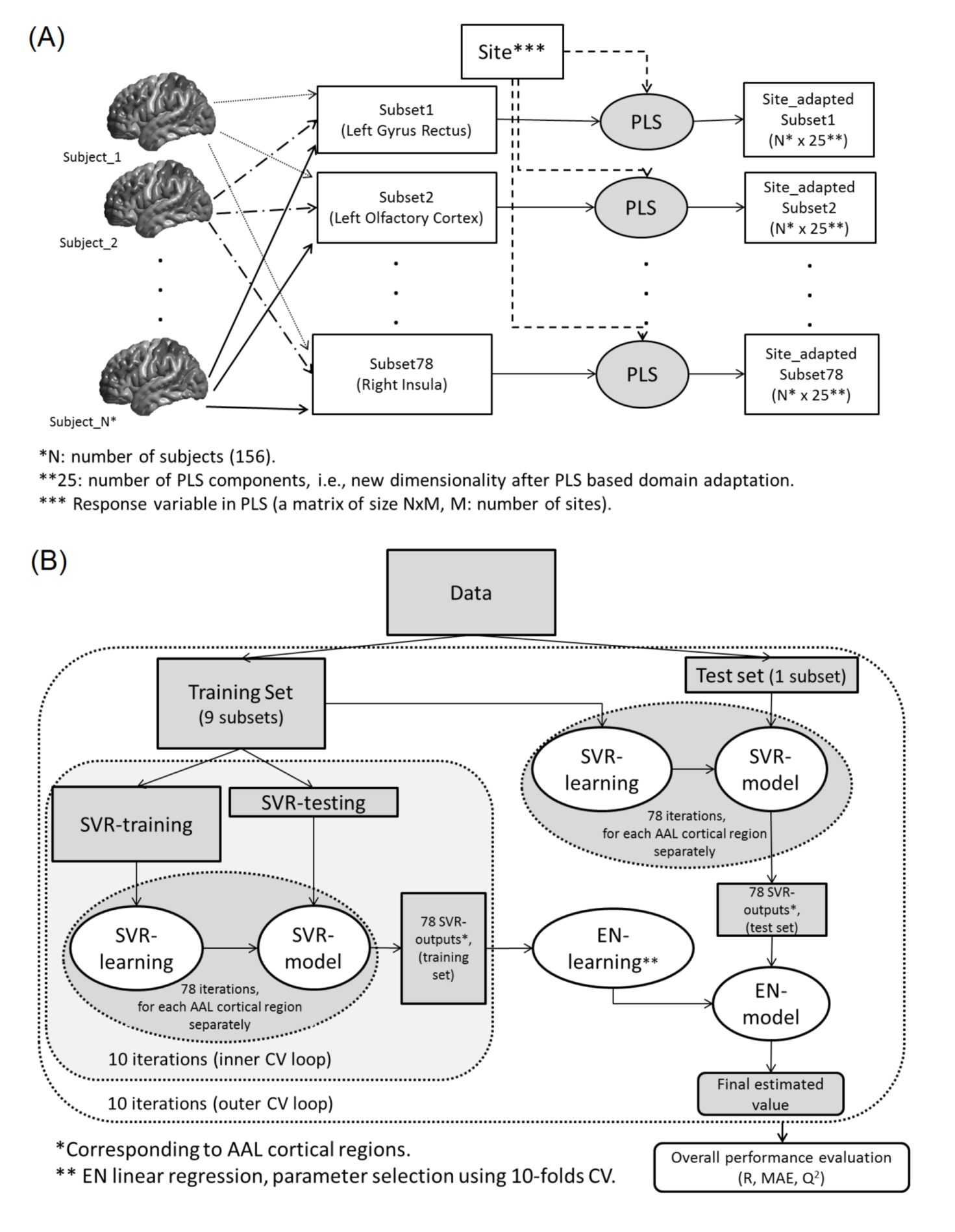
Workflow of the proposed method for estimating severity score in ASD subjects. A) The PLS based domain adaptation stage and B) the learning stage.

In the learning stage, first, we applied SVR in each (site-adapted, after domain adaptation) subset separately, with the severity score as the response variable (Section 2.7). This resulted in 78 outputs, each of them estimating the severity score based on only one AAL brain region. In order to combine the results from different brain regions, we concatenated these 78 outputs from SVR to form a new dataset. The resultant dataset has dimensionality 78, from 78 SVR outputs. Finally, we applied elastic-net penalized linear regression on the new set to obtain the final estimated severity score (Figure 1 B; Section 2.8).

### 2.6. Partial Least squares domain adaptation

As our data are from 4 different sites, our purpose is to identify a feature space where the data from different sites have similar distributions. We propose to achieve this by using Partial Least Squares (PLS) in order to identify a new low dimensional feature space that would only contain such cortical thickness information that is maximally invariant between the acquisition sites. PLS is a linear feature transformation method for modeling relations between sets of observed variables. Similarly to principal component analysis (PCA), PLS constructs new predictor variables, i.e., latent variables, as linear combinations of the original predictor variables; regional cortical thickness values in this case. The difference between PCA and PLS is that PLS considers response variables, sites in our case, for constructing latent variables while PCA considers only the predictor variables. In particular, PLS tries to discover the relation between the predictor variables **X** and response variables **Y** by determining the multidimensional direction in the **X** space with the maximum multidimensional variance direction in the **Y** space.

We denote a regional subset of the cortical thickness values by **X** ∈ ℝ*^N×D^*, where *N* is the number of subjects and *D* is the number of cortical thickness measures in the corresponding subset. *D* varied from 114 (olfactory cortices) to 2218 (middle frontal gyri). The same process is applied to each of the 78 cortical regions; we drop the sub-section index for clarity. The response variable representing the site information is **Y** = {*Y*_1,1_,…,*Y_N,M_*}, where *M* is the number of sites. *Y_n,m_* is 1 if subject *n* belongs to site *m* and otherwise it is 0. PLS assumes the following relationships between **X** and **Y**:

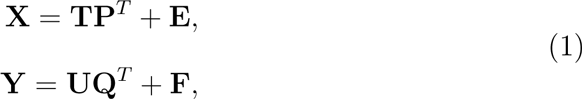

where the latent variables corresponding to **X** and **Y** are stored in **T** and **U** matrices, respectively; **P** and **Q** are loading matrices and **E** and **F** are error terms. In particular, the *N* × *K* matrix **T** = [**t**_*1_,…, **t**_\**K*_] = [**t**_1_,…,**t***_N_*]*^T^*, where *K* denotes the number of PLS components, provides projections of cortical thickness values that we are going to use to predict severity scores. The decompositions of **X** and **Y** are computed by iterative application of the singular value decomposition (SVD)(Abdi, 2007; de Leeuw, 2007) in such a way that in each iteration the covariance between **T** and **U** is maximized. That is, in each iteration, PLS tries to find weight vectors **w_i_**, **c_i_** so that

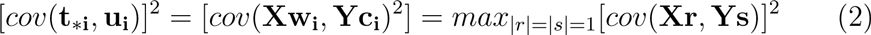

where *cov*(**t_*i_**, **u_i_**) = **t_*i_***^T^* **u_i_**/*N* is the covariance between latent variables corresponding to the cortical thickness and the site information (Rosipal and Krämer, 2006). For the computation of PLS, we use the SIMPLS algorithm (De Jong, 1993) that yields cortical thickness projections **t**_\**i*_ directly as linear combinations **Xw**_*i*_ and, importantly, constraints any **t**_*\*i*_ and **t**_*\*j*_ to be orthogonal. The idea is that the first few **t**_*\*i*_ (*i* < *V*) encode the site related information and then the later **t**_*\*i*_ (*i* ≥ *V*) contain site invariant information; note that *V* may have the value of 1. In this reasoning, we utilize the connection between the PLS and the Fisher’s discriminant analysis (Rosipal and Krämer, 2006). We leave it to the machine learning algorithm to discard the first components that may be useless for the severity score prediction and keep all the PLS components.

We note that PCA, but not PLS, has previously been used for unsupervised domain adaptation as a baseline method for the applications of object recognition and sentiment analysis (Shi and Sha, 2012), where all data from both source and target domain were projected into PCA direction computed from the data in the target domain. In Shi and Sha (2012) the model was trained on a data from the single source domain and tested on data from the target domain while we consider the multiple source domain adaptation.

We have additionally developed and tested an inductive version of the algorithm which comes with certain disadvantages compared to the transductive version. These and experimental results with the inductive algorithm are discussed in the Section 4 of the supplement.

### 2.7. Support Vector Regression

After PLS analysis on each of the 78 regional subsets of cortical thickness measures, we have 78 matrices **T***_ℓ_*, *ℓ* = {1,…, 78} of the site adapted cortical thickness coefficients corresponding to the 78 cortical regions. To derive a prediction of the severity score based on a single cortical region, we apply support vector regression (SVR). Again, the process is done independently for each region and we drop indexes pertaining to the regions for clarity.

Support vector machines (SVM) were first introduced (Cortes and Vapnik, 1995; Boser et al., 1992; Vapnik and Vapnik, 1998) as a pattern recognition method representing decision boundary between samples from two different classes in such a way that the margin (the distance) between the decision boundary and the closest training sample to it is maximized. SVM transforms the training data from the original space into a high dimensional feature space via a kernel-induced mapping function, and then the separating hyperplane is computed in this new feature space.

Support vector machines can also be applied to regression problems when the response variable is a real-valued number, resulting in support vector regression (SVR). To achieve the maximal margin property in a regression problem, Vapnik (1995) proposed the ε-SVR algorithm by devising the ε-insensitive loss function. In SVR, a specific value is determined as ε in the loss function, after which the task is to fit a regression line surrounded by a tube with radius ε to the data. The data points inside the tube are not considered in determining the regression line and only the data points lying on the edges or outside the tube, i.e. support vectors, affect the course of the regression line.

SVR approximates a severity score by a nonlinear function described by the weight vector 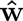 and the bias 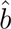 so that

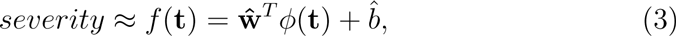

where **t** is a vector of the regional site adapted cortical thickness (CT) coefficients for a subject, ϕ is a non-linear mapping and the response variable is the corresponding severity score. SVR handles the nonlinearity via the kernel trick. A high (or infinite) dimensional dot product 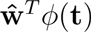 can be computed as a sum of dot products implicitly described in the input space with the original dimensionality 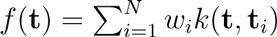, where *k* is the kernel function, **t**_*i*_ are the site adapted CT coefficients for the training subject *i* and *w_i_* are the parameters to be solved by the SVR algorithm. The kernel-trick makes otherwise intractable computations feasible and *ϕ* and 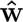 do not need to be explicitly defined. In this work, we adopted the radial basis function kernel (RBF) *k*(**x**, **y**) = exp(−γ||x − y||^2^) and set γ to its default value 1/*K*, where *K* = 25 is the number of PLS components. The RBF kernel is the most widely used kernel function in nonlinear SVR. For solving the SVR parameters *w_i_*, 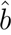, we used ν-SVR (Schölkopf et al., 2000). This is a re-parametrization of the original soft-margin *ε*-SVR algorithm (Cortes and Vapnik, 1995) allowing automatic tuning of *ϵ* by introducing an additional parameter *ν* (Smola and Schölkopf, 2004). The *ν*-SVR aims to solve the following optimization problem:

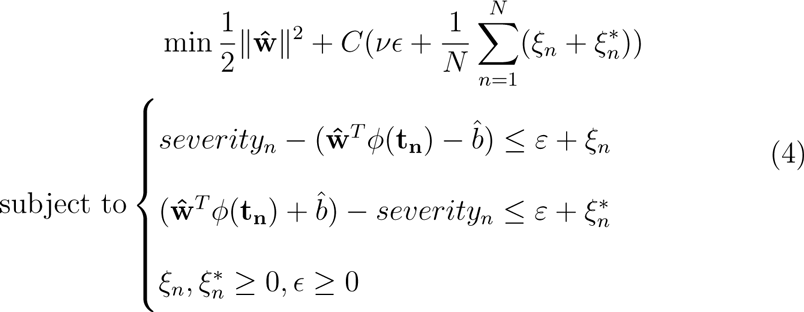

This allows for training errors exceeding *ϵ* by introducing slack variables 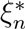. The overfitting is prevented by the regularization term 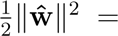 = 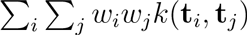 and the tradeoff between the close fit to the data and regularization is controlled by the parameter *C*.

We re-iterate that the purpose of this step is to determine a predictive severity score for each subject based on each cortical region. This step was repeated for each brain region separately, which resulted in 78 single scores for each subject, each of them predicting severity score based on one cortical region.

### 2.8. Penalized Linear Regression

From the SVR, we have a predicted severity score 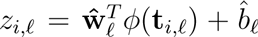 for a subject *i* and region *ℓ*. For each subject *i*, we concatenate the regional predictions into a 78-element vector **z**_*i*_. In order to integrate the predicted severity scores derived from different brain regions, we used least squares linear regression (LR) with elastic net penalty. The elastic net penalty is a combination of ridge and lasso penalties (Zou and Hastie, 2005) that has two important advantages in our case: 1) it allows for variable selection, meaning that the regions with low predictability are dropped from the model and 2) it possesses the grouping effect meaning that the regions with similar predictions receive similar weights in the final model. These two properties improve the interpretability and stability of the elastic-net penalized models. The LR model is formalized as:

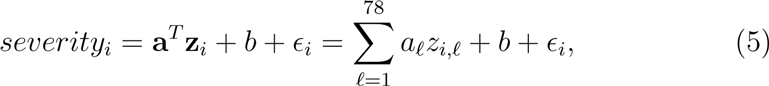

where *i* refers to a subject, **a** = [*a*_1_,…, *a*_78_]*^T^* and *b* are the model parameters and *ϵ_i_* is the error term. Adding the elastic net penalty, the model is solved by minimizing the following elastic net cost function:

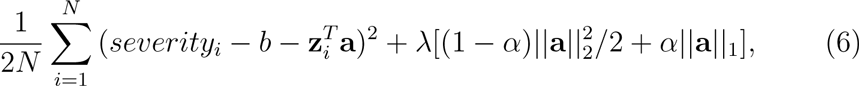

where *N* is the number of training samples, λ is the complexity parameter found by cross-validation, *α* ∈ [0, 1] defines the compromise between ridge 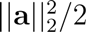 and lasso penalties ||**a**||_1_, and || · ||_1_ denotes the L1-norm. Here, we selected *α* = 0.5 to give equal weights for the ridge and lasso penalties.

### 2.9. Implementation and validation

It is imperative to avoid using the test subjects’ severity scores for training the model as this would result positively biased estimates of the prediction accuracy. For dividing data into two training (SVR-training and SVR-test) and test sets, we used two nested and stratified cross-validation loops (10folds for each loop) except for site-based testing where the outer loop was leave-one-site-out loop. In the inner CV loop, the SVR-train set was used to train the SVRs and the SVR-test set was used for constructing regional predictions *z_i,ℓ_* for every training subject; we did not use the same dataset both for learning the SVR and computing regional predictions to avoid overfitting. The training set (union of SVR-training and SVR-test) was used to train the Elastic-net regression model. We re-divided the training set into 10-folds for finding the optimal λ for the model. Test data were used only for evaluating the model. The performance of the model was then evaluated based on the (cross-validated) Pearson correlation coefficient (*R*), mean absolute error (*MAE*) and the coefficient of determination^3^(*Q*^2^) between estimated and true severity score in test set. The reported results are averages over 100 nested 10-fold CV runs in order to minimize the effect of the random variation. Three different metrics are reported, because these each provide complementary information. Cross-validated *R* is simple to interpret, but it can hide the bias in the predictions, which are made apparent by *Q*^2^-value. MAE provides the prediction errors in the equal scale with the original scale of the severity scores. Prior to each step, both the predictor variables and response variable were normalized to have zero mean and unit variance, except in domain adaptation step in which the data are centralized/normalized by default. To compare the performance of two learning algorithms, we computed a p-value for the 100 correlation scores with a permutation test. For computing p-values associated with the null hypothesis that the correlation coefficient between the observed and predicted values is zero, we used a permutation test (Anderson and Robinson, 2001) and for computing the 95% confidence interval of the correlation coefficient we used bootstrap on the run with the median correlation score across 100 cross-validation iterations.

PLS was computed by the PLSREGRESS functions in MATLAB software with a fixed number of components. The SVR training was implemented using LIBSVM (Chang and Lin, 2011). The parameters in SVR, namely *C*(the soft margin parameter) and *γ* (parameter for RBF kernel function), were set to their default values (*C* = 1, *ν* = 0.5, *γ* = 1/*F*, where *F* is the number of features here equaling to *K* = 25). Since the cortical thickness measures were divided into 78 subsets and both PLS and SVR were computed in each subset separately, tuning the method parameters, inside a nested cross-validation loop, was impractical. Therefore, we used fixed number of components in PLS and the default parameters of the SVR across all subsets. The fixed number of PLS components in the proposed method was 25, selected by initial experiments among the candidate set {5,10,15, 20, 25, 30}.

The implementation of elastic-net penalized linear regression was done by using the GLMNET library (Qian et al., 2013) and the regularization parameter *λ* was selected using 10-folds CV in the training data. Note that the penalized LR was done only once in the outputs of SVR from different brain regions and hence tuning the regularization parameter using CV was easily feasible.

## 3. Results

The average cross-validated correlation *R* between the estimated and observed severity scores among 100 distinct 10-fold CV iterations was 0.51 (standard deviation 0.04, range from 0.39 to 0.63, *p* < 0.0001), the average mean absolute error (MAE) was 1.36 (standard deviation 0.05, range from 1.25 to 1.51) and the average coefficient of determination *Q*^2^ was 0.26 (standard deviation 0.045, range from 0.13 to 0.39). These values indicated that the proposed approach was able to provide information about the severity of the disease based on structural information of the brain in ASD patients. Particularly, we note that the union of 95% confidence intervals (CIs) of *R* for individual runs was [0.25, 0.72], where CIs were computed based on the Fisher’s r-to-Z transform, and the lower limit of the worst 95% CI of *R* was clearly positive. The box-plots of the correlation scores and MAEs are available in Fig. 2 and the scatter plot of the estimated and observed severity scores of the CV run with the median *R* is shown in the upper left panel of Fig. 3. We note that validation accuracy was almost the same (the average *R* was 0.49 or 0.50 depending on whether module information was used) when predicting raw ADOS scores instead of the proxy severity scores. The validation results concerning the prediction of the raw ADOS scores are presented in the Supplementary figures 2, 3, and 4.

**Figure 2:**
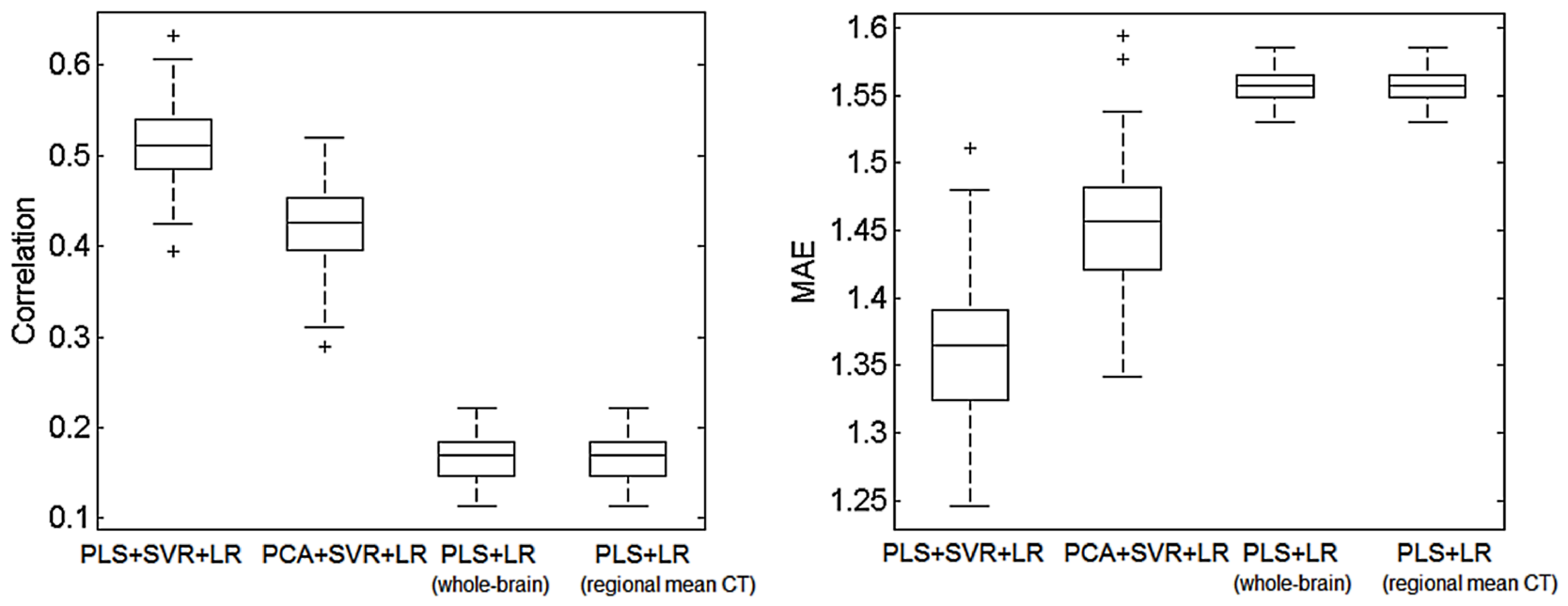
Box plots for correlation score and mean absolute error within the 100 computation runs of the proposed approach (PLS + SVR + LR), substituting PLS based domain adaptation by PCA (PCA + SVR + LR) and without the SVR step (PLS + LR). PLS + LR (whole brain) refers to the approach where all 81924 vertices were used as the input to PLS stage and PLS + LR (regional mean CT) refers to the approach where the regionally averaged thickness values were used as the input for the PLS; see the text for details. On each box, the central mark is the median, the edges of the box are the 25th and 75th percentiles, the whiskers extend to the most extreme data points not considered outliers, and outliers are plotted with a +.

**Figure 3:**
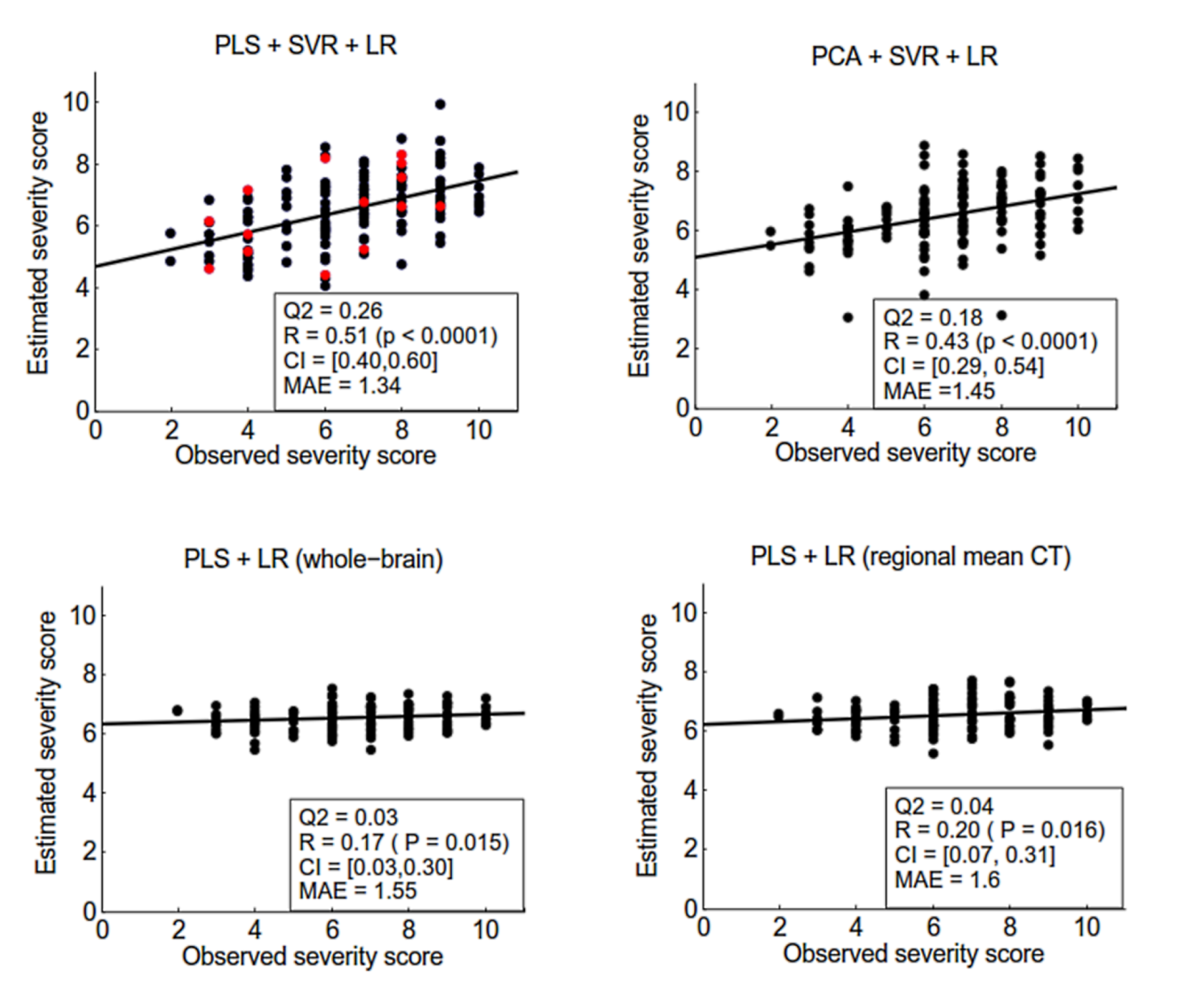
Scatter plots of the estimated severity score vs. observed severity score for the proposed method (PLS + SVR + LR), without PLS based domain adaptation (PCA + SVR + LR), and without the SVR step (PLS + LR). See the text and Figure 2 for details. The scatter plots are from a cross-validation run with the median correlation within 100 computation times. In the panel corresponding to PLS + SVR + LR, data corresponding to female subjects is shown in red color in order to ensure that they did not act as outliers.

**Figure 4:**
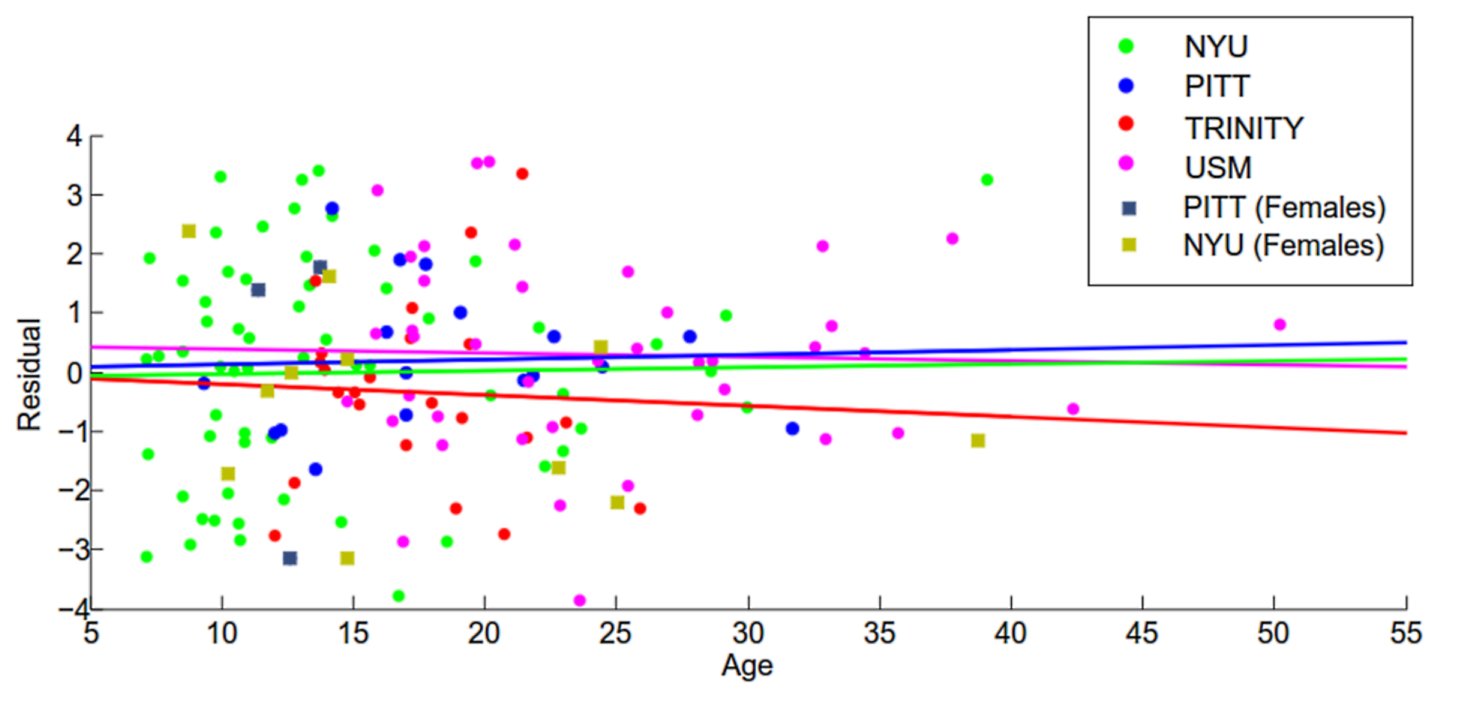
Scatter plot of prediction residual vs. age for the proposed method (PLS + SVR + LR) with a cross-validation run with the median correlation score within 100 computation runs. A fitted line is added for the residuals of each site. There was no significant difference within the slopes of fitted lines (*p* > 0.5) and the slopes of all fitted lines are non-significant (*p* > 0.5). Female subjects are plotted with a different color than the male subjects, however, the regression lines were fitted considering both genders together.

For evaluation of the effectiveness of each stage (PLS, SVR, Elastic-net LR) of the proposed approach, we performed experiments by excluding each stage of the method separately and comparing the accuracy of the predictions obtained this way to the accuracy of the predictions of the complete method.

To evaluate the PLS based domain adaptation stage, we repeated the experiments with the same procedure, except that we replaced PLS by PCA which can be thought as an unsupervised dimensionality reduction method equivalent to PLS but not utilizing the information about the acquisition site. In other words, by using PCA, a common feature space was determined for all data from different sites without considering the site information. The PCA was applied in the transductive setting as The optimal number of PCA components used (20) was selected with the same procedure as the number of PLS components (see Section 2.9). When the PLS-based domain adaptation was substituted by PCA, the average correlation score (among 100 different runs) dropped from 0.51 to 0.42 (*p* < 0.0001 for correlation decrease), the average MAE increased from 1.36 to 1.45 and the average *Q*^2^ dropped from 0.26 to 0.17. Since both PCA and PLS project data into a new feature space, we omitted this feature transformation step to see the effect of image acquisition differences between sites on the performance of the model. When the the feature transformation step was omitted, the average correlation score (only 5 CV runs were done) decreased to 0.16, the average MAE increased to 1.65 and the average *Q*^2^ dropped to −0.07. Thus, the feature transformations were useful.

To validate the SVR step, we performed two experiments. First, we estimated severity score by applying elastic-net penalized regression directly on the site adapted thickness values, i.e., retaining PLS-based domain adaptation step but performing it to the 81924 thickness values without dividing them to regional subsets and not performing the nonlinear SVR (PLS + LR (whole brain)). By eliminating the SVR step, the average correlation score decreased to 0.17 (*p* < 0.0001 for the correlation decrease), the average MAE increased to 1.56 and the average *Q*^2^ decreased to 0.03. Second, we averaged the cortical thickness values within each AAL region, performed the PLS based domain adaptation on these 78 regional mean cortical thickness measures and used the Elastic net penalized LR to predict severity scores based on the resulting PLS components (PLS + LR (regional mean CT)). The average correlation score decreased to 0.20, the MAE increased to 1.55 and the *Q*^2^ decreased to 0.04. Again, the optimal number of the PLS components (5) was selected by the same procedure as for the complete method (see Section 2.9). We also repeated the experiments (only 5 CV runs were done) by omitting the PLS step and applying elastic net penalized LR on regional mean of cortical thickness to predict severity scores. This experiment yielded the average correlation score of 0.05, the average MAE of 1.60 and the average *Q*^2^ of −0.02 and it appeared that the severity cannot be estimated based on the regional cortical thickness values.

Figure 2 shows box plots for the *R* and *MAE* for different experiments across 100 computation runs. It can be observed that the regional SVR had the largest effect on the performance of the method. The performance of the method was not good when excluding this step despite that PLS based domain adaptation was used. Figure 2 also illustrates that the PLS based domain adaptation step led to markedly improved predictions when coupled with the regional SVR. Figure 3 shows the scatter plot between estimated and observed severity scores (of the median correlation within 100 computation times). According to these plots, the severity scores with very high or very low values were the most difficult to estimate as most of the observed severity scores were located within the range from 4 to 9. Also, as shown in the upper left panel of Fig. 3, the few females in the sample did not act as outliers. Figure 4 illustrates the effect of age on the estimated severity scores for the proposed approach. As it can be seen from the Figure 4, there is no effect of age on the residuals and there is no significant difference within the residuals of different sites. The results of an experiment performed with a more narrow age range are reported in Section 5 of the supplement.

Figure 5 shows the importance of top 24 brain regions identified by average magnitude of the regression coefficients in the penalized LR, i.e., the final step of the proposed approach, within 100 computation times of 10 fold CV. The visualization of these regions is provided in Figure 6. Since we standardized the data before applying LR, the absolute value of each regression coefficient provides the importance of corresponding predictor in the model and therefore we could compute the importance of each brain region based on the magnitude of the regression coefficients.

We studied the effect of acquisition site on the performance of the proposed method. To address this issue, a “site-wise” cross-validation analysis was performed. To be more specific, a 4-fold leave-one-site-out CV was performed in such a way that the data from each site was in its own fold and the method was trained using data from 3 sites and tested in the remaining site. The results are listed in the Table 2. Figure 7 shows the scatter plot between estimated and observed severity scores (of the median correlation within 100 computation times) for each site. The prediction accuracy of the site PITT was comparable with that of the standard 10 fold CV, but the prediction accuracy in the other sites decreased markedly from that of the standard 10 fold CV. These results suggest that utilizing some samples from the same site as the test sample in the learning procedure might improve notably the prediction accuracy. One possible explanation for this result is obviously the decreased number of training subjects available for the method training, especially in the case of NYU and USM sites, which contained the largest number of subjects (NYU 72 of 156 subjects and UsM 41 of 156 subjects, see Table 1). Also, *Q*^2^ scores for TRINITY and USM sites were strongly negative indicating that the severity scores predicted from the data of the other sites were biased. One reason for the bias can be explained when examining the average observed severity scores from each site (NYU: 6.3; PITT: 6.7; TRINITY: 5.7; USM: 7.4). The average severity score of TRINITY was lower than the average of the other sites and the average severity score of USM was higher than the average of the other sites while the penalized regression creates shrinkage towards the average severity score (see Zou and Hastie (2005)) and thus could produce biased severity predictions for the two sites. We note that the domain adaptation method of this article cannot correct for possible site differences in administering the ADOS tests as it is blind to severity scores.

**Table 2:** The results of “site-wise” based cross-validation. The reported results are the average between 100 computation times.

| Site | Correlation | MAE | $Q^2$ |
| --- | --- | --- | --- |
| NYU | 0.22 | 1.57 | -0.04 |
| PITT | 0.56 | 1.08 | 0.22 |
| TRINITY | 0.15 | 1.59 | -0.25 |
| USM | 0.24 | 1.44 | -0.29 |

**Figure 5:**
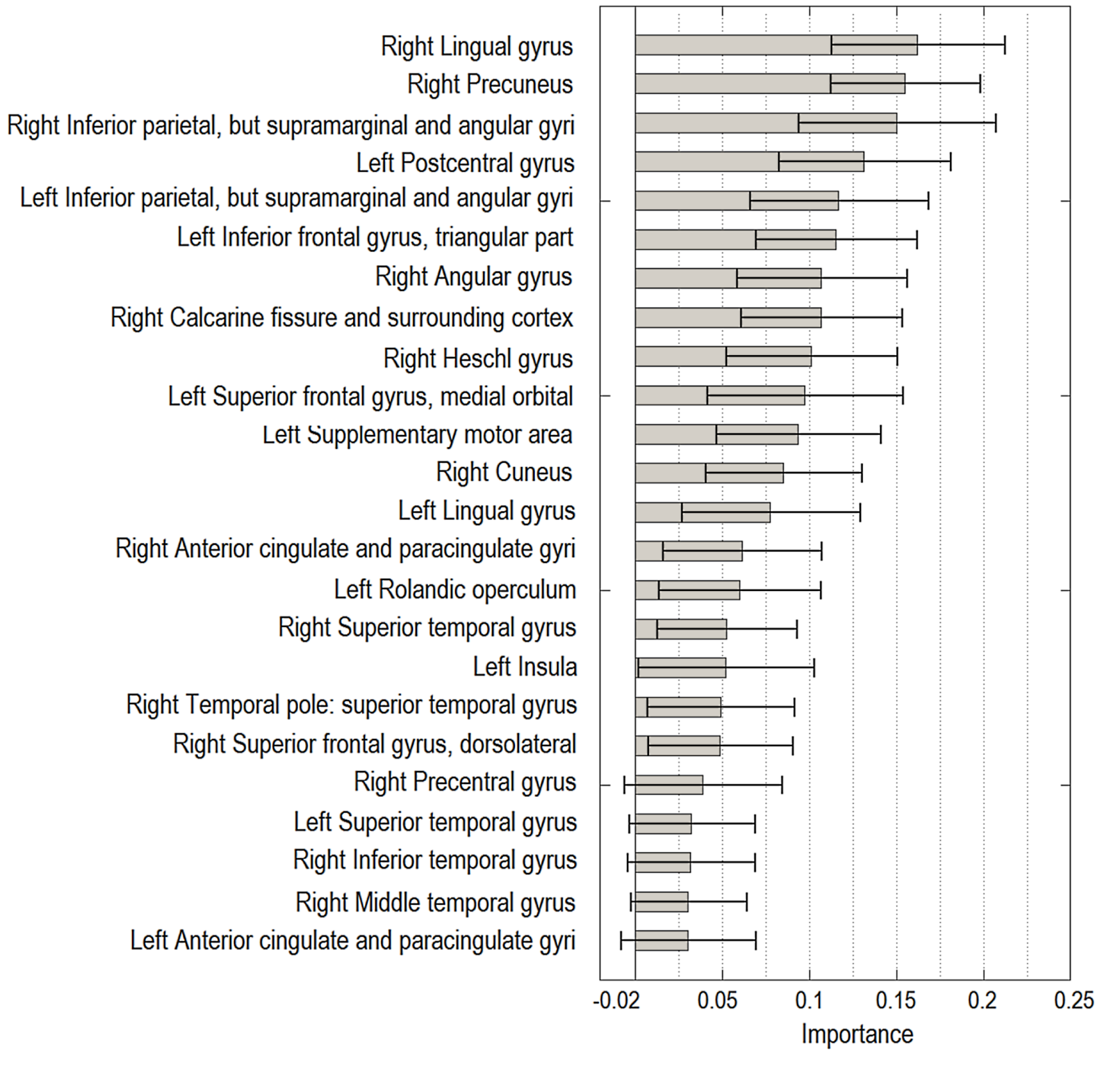
The importance of the top predictors for estimating severity score in ASD subjects. The ranking is based on the average magnitude of standardized regression coefficients across 100 cross-validation runs. The gray bars display the average magnitude and the error bars (in black) of the length equal to twice the standard deviation of the magnitude. Predictors with the average magnitude higher than 0.03 are included. For the importance of other regional predictors, see Fig. 6.

**Figure 6:**
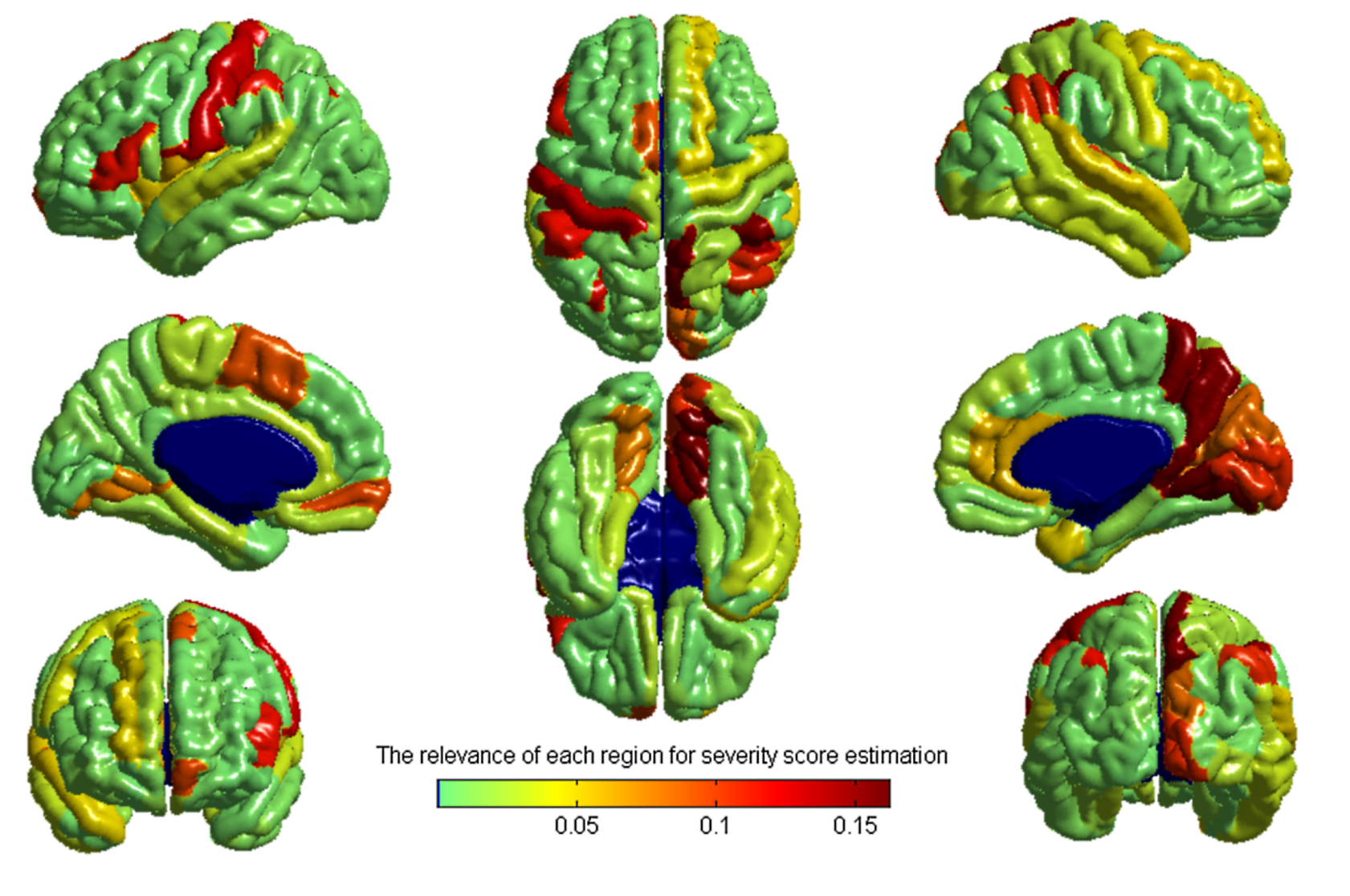
The importance of each cortical region in the estimation of severity score using the proposed approach. The importances are the average magnitudes of the standardized regression coefficients from the Elastic-net penalized regression across 100 cross validation runs.

We experimented with the method by training and testing with single site data, that is, we trained four different prediction models and tested them with the data from the same site in the nested cross-validation framework. The average cross-validated correlation *R* within ten 10-fold CV runs was the largest for the site USM (average correlation score *R*(*USM*) was 0.22) and for the three other sites the average correlation score was close to or below zero (*R*(*NYU*) = −0.05, *R*(*PITT*) = 0.01, *R*(*TRINITTY*) = −0.28). These results clearly suggested the utility of having a larger number of subjects at the expense of having to deal with multi-site data. We still point out that the variance of cross-validated performance measures was inflated due to small sample sizes and the sample sizes for PITT and TRINITY are too small for adequate error estimation. In particular, the clearly negative *R* for the site TRINITY, with the smallest sample size, could be attributed to the small sample size that, for example, considerably decreased the stability of the inner CV and led to the selection of poor models.

Since certain cognitive functions are lateralized (Hugdahl, 2005), we performed the experiments within right and left hemispheres separately to study the relative relevance of each hemisphere in estimating the severity score. The results revealed greater relevance of right hemisphere in estimating the severity score compared to left hemisphere. Using only the cortical thickness measures belonging to the right hemisphere yielded the average correlation score of 0.46, the average MAE of 1.41 and the average *Q*^2^ of 0.20. The measures in the left hemisphere produced significantly lower average correlation score of 0.28 (*p* < 0.0001), the average MAE of 1.53 and the average *Q*^2^ of 0.05. These results support the findings of Torgerson et al. (2015) that indicated higher relevance of regions and connections of the right hemisphere compared to the left hemisphere in predicting ASD severity based on ADOS score. While using cortical thickness measurements from only the right hemisphere led to accurate severity score estimates, combining cortical thickness measurements from both right and left hemispheres still led to a better performance (*p* < 0.0001). This can be also seen in Fig. 5 where among the most important brain regions for the model there are regions from both hemispheres, although, the best predictors were located in the right hemisphere.

**Figure 7.**
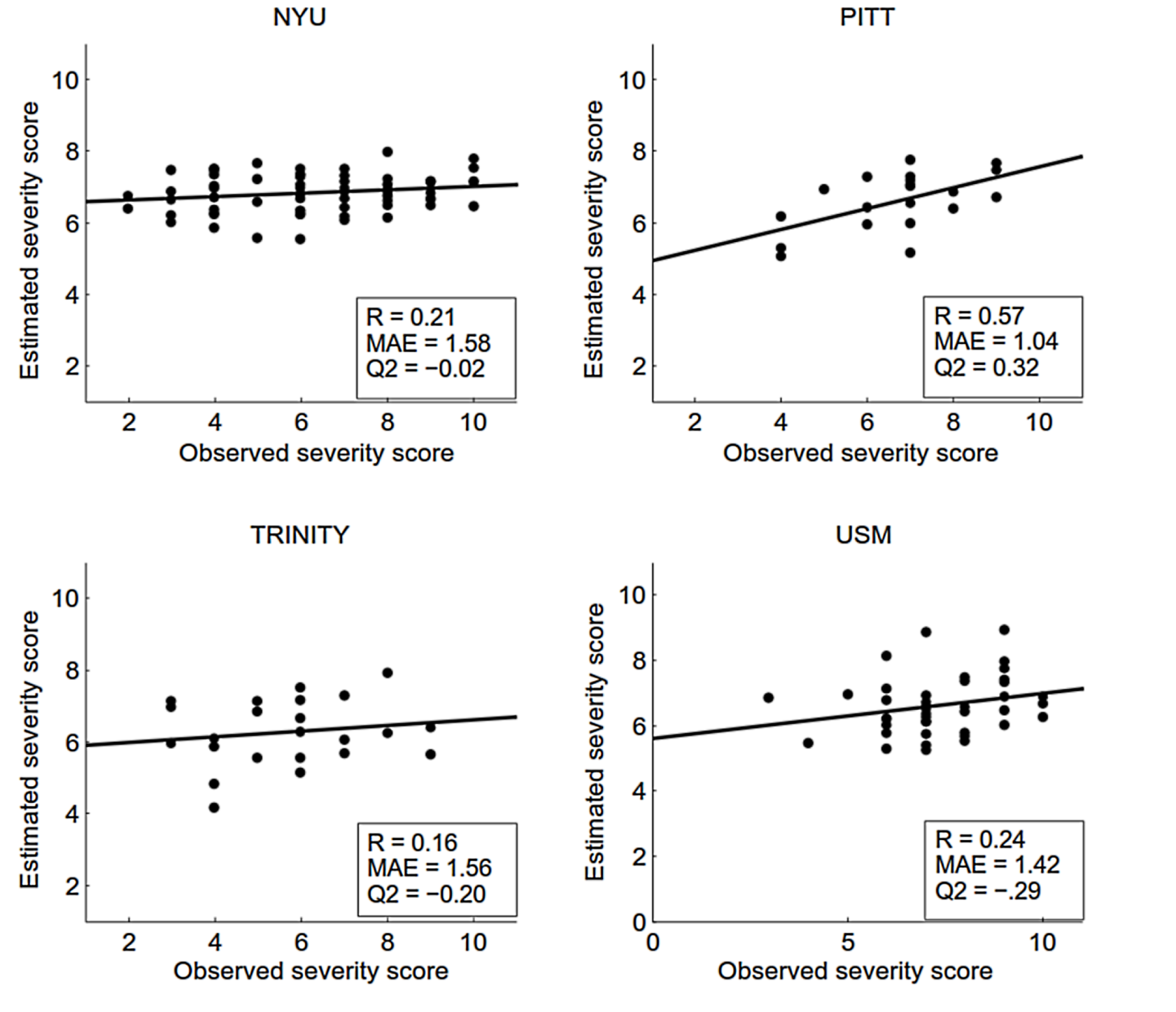
Scatter plots of the estimated severity score vs. observed severity score for the proposed method for each site separately. The scatter plot for different sites are from a cross-validation run with the median correlation within 100 computation times.

In order to demonstrate the suitability of SVR (with an RBF kernel) for designing regional models in the proposed approach, we replaced it with different linear models (elastic net LR, relevance vector regression (RVR) and SVR with linear kernel) for predicting severity scores. Replacing the non-linear SVR with the linear alternatives led to a marked performance decrease. The correlation score averaged over 10 CV runs dropped to 0.32 when using linear SVR, 0.28 when using linear RVR and 0.13 when using elastic net LR. The elastic net LR was selected as the learner for the last step to obtain a model that is easy to interpret and we did not test other learners for this stage. As explained in Section 2.8, the elastic net LR provides spatially sparse model by simultaneously performing variable selection and model estimation and, furthermore, it possesses so called grouping effect meaning that correlated predictors are selected simultaneously (Zou and Hastie, 2005).

**Figure 8.**
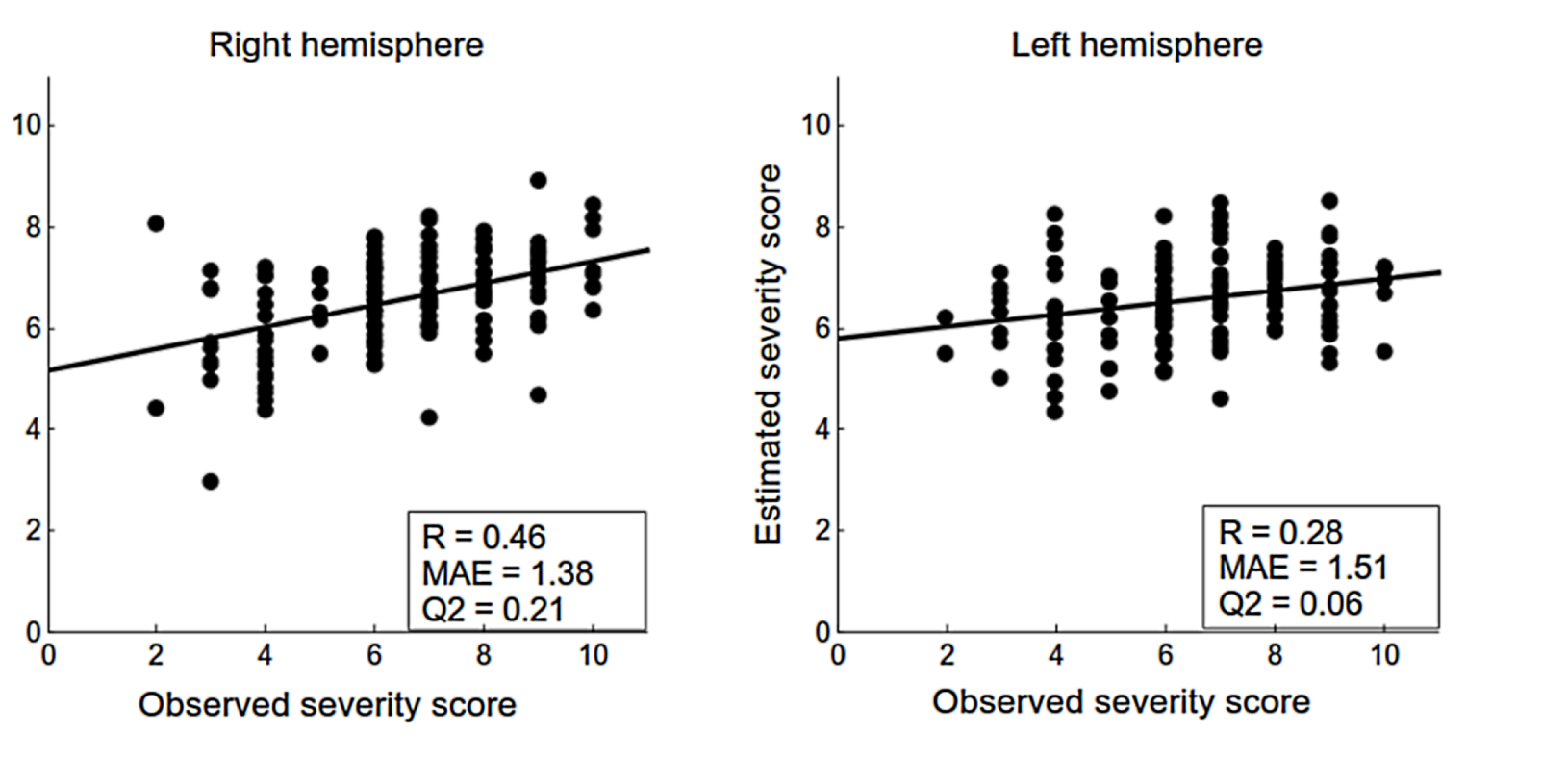
Scatter plots of the estimated severity score vs. observed severity score for the proposed method for each brain hemisphere separately. The scatter plots are from a cross-validation run with the median correlation within 100 computation times.

## Discussion

The objective of the current study was to devise methods to overcome the issues associated with multi-site, multi-protocol data in order to take advantage of the increased sample sizes provided by such agglomerative data to better predict behavioural outcomes from brain structure. We explored this problem using data from four sites from the ABIDE dataset, and used cortical thickness to predict ADOS-based ASD symptom severity. We developed a novel two-stage approach consisting of a domain adaptation stage that uses partial least squares regression with site as a response variable, and a learning stage which utilizes a combination of support vector regression and linear regression. We evaluated the reliability of the method by comparison with variations without domain adaptation, or without support vector regression. The proposed two-stage method performed markedly better than the alternatives, and resulted in a cross-validated correlation score that was much higher than for any of the sites alone, and considerably higher than has previously been reported in the literature for multisite data (Sato et al., 2013).

Recent studies on multisite classification of autism using ABIDE data have shown poor accuracy in classification of ASD versus TD subjects (Nielsen et al., 2013; Haar et al., 2016). The study by Nielsen et al. (2013) showed that classification rate was much lower in a multisite dataset than for single site data. The effect of scanner variation in multisite analyses of cortical thickness abnormalities in ASD patients was also studied by Auzias et al.(2014, 2016). They showed that scanner variation is a significant confounding factor, which is distributed across the cortical surface and reaches its peaks in the frontal region. Thus, the effect of acquisition site on the basic image properties might be a possible reason for the poor classification accuracy in the studies by Nielsen et al. (2013) and Haar et al. (2016), as well as for the inconsistencies on the reported results from different studies, especially in the context of abnormalities in cortical thickness measurements (Raznahan et al., 2013; Hadjikhani et al., 2006).

In the current study, we used PLS based domain adaptation in order to maximize the consistency of the imaging measures over the multiple scanners/protocols before assessing ASD pathology. Unlike previous approaches, such as PCA, in which site/scanner are treated as any other nuisance variable, the PLS based domain adaptation established a feature space where the data from multiple sites/scanners have similar distributions. Accommodating multiple sites/scanners in such a way resulted in significantly improved performance (Figs. 2 and 3), indicating the power of our PLS based domain adaptation approach for dealing with multi-site data. While our domain adaptation method can correct for differences in imaging data between sites, it cannot correct for possible site differences in administering the ADOS tests (due to inter-examiner differences in the administration and scoring of the tests) as it is blind to severity scores. Also, the domain adaptation method searches for consistent data projections across sites and tries to divide the thickness data in the orthogonal site-specific and site independent components. Therefore, it has no control what is the cause for the site-specificity of the data later left out by the SVR (scanner differences, different subject characteristics, or interactions of the two, which are all characteristic to neuroimaging data agglomeration efforts). The method needs a certain number of subjects for each site and we have no clear answer what this number should be. We also hypothesize that the necessary number of subjects per site increases with the number of different sites, as the site adaptation problem becomes harder as more sites need to be accommodated in a common feature space. More specifically, the complexity of the PLS based domain adaptation step increases when more sites are added due to the increase of the dimensionality of the response variable. Including those 4 sites which had at least 20 ASD subjects with severity scores available led to promising results in this study and the requirement of having this number of subjects per site does not limit the foreseeable applications of the method.

The subjects ranged in age from 8 to 40 years, and the age is known to influence cortical thickness in ASD (Doyle-Thomas et al., 2013). Note that while the age influences cortical thickness, it can be assumed independent of the severity score due to the calibration, and, therefore, it acts as a source of nuisance variability for the prediction (similarly to so called suppressor variables in the ordinary linear regression (Friedman and Wall, 2005)). Therefore, the age effects on cortical thickness do not artificially increase the cross validated performance measures, but accounting for them could improve the predictions and we tried to incorporate age information in the learning process, in order to improve disease severity predictions. However, the experiments with multiple methods were unsuccessful with the best results reached by including the subject age in the domain adaptation step so that the response variable in PLS was constructed based on subject’s site and age information. However, by doing this the performance of the model dropped considerably (cross-validated R was 0.44), technically probably due to increase of the complexity of the domain adaptation. Linearly regressing out the age information, that is a widely used in dementia related machine learning applications (Klöppel et al., 2015) and has often improved the predictions (Tohka et al., 2016), did not work here (*R* was 0.42 when the age was regressed out vertex wise before the domain adaptation step and *R* was 0.30 when the age was regressed out component-wise after the domain adaptation). We speculate that these results are due to 1) less pronounced age related cortical thickness changes in autistic subjects than those of normal controls (Doyle-Thomas et al., 2013); 2) strong variation in the age related change according to the disease severity, which undermines the suitability of severity score independent age corrections; and 3) since the age is one of the probable sources of the data heterogeneity, possibly projecting the data in the new space manages to separate some of age effects into their own components aiding machine learning algorithm to handle the nuisance variability caused by age. Finally, results with the data set with a more restricted age range are reported in the supplement (Section 5) suggesting that, for our method, it is more important to have a larger number of training subjects than to try to balance the subject demographics across the sites.

Haar et al. (2016) suggested that their poor decoding accuracy for classification of multisite ABIDE data was not only because of between-site variation, but also weak anatomical abnormalities in the ASD pathology which offer very limited diagnostic value. Substantial variability within each diagnostic group complicates classification, hence our decision to predict symptom severity from neuroimaging measures. The prediction of raw ADOS scores based on MRI and cortical thickness was previously investigated by Sato et al. (2013). They predicted ADOS from MRI based inter-regional thickness correlations with SVR as the machine learning method. The method yielded a cross-validated Spearman correlation of 0.36 with a dataset consisting of MRIs of 82 autistic patients acquired at three different sites with a standardized protocol. To compare our results to theirs, we calculated the cross-validated Spearman correlation between the estimated and observed severity scores, which was 0.51. The higher correlation value that we obtained must be understood in the context of the following differences between our study and that performed by Sato et al. (2013). First, our data are from 4 different sites without any standardization protocol, so the between-site variation was an additional challenge in the current work. Second, Sato et al. (2013) used inter-regional thickness correlation for estimation ADOS score, instead, we determined a predictive score for each distinct brain region and then combined them via a linear regression model to estimate severity score. Third, we used severity score instead of using raw ADOS score. Lastly, our method was evaluated with almost double the sample size (156 subjects).

In addition to the PLS-based domain adaptation, the other novel technical characteristic of the proposed method was our treatment of the whole-brain problem of prediction as a set of regional problems of prediction. We divided the cortical thickness measures into regional subsets, determined a predictive score for each region separately, and then combined the regional scores into a whole brain measure of disease severity. This enabled us to divide the problem into several sub-problems with lower complexity while better retaining the original spatial resolution of the thickness measures. We hypothesized that both of these properties are important for successful predictions: Khundrakpam et al. (2015) have previously demonstrated that a fine parcellation of the cortical thickness measures was advantageous for age estimation within healthy children. However, increasing spatial resolution results in higher dimensionality, which increases the complexity of the model. Specifically, in the domain adaptation stage, finding a low dimensional site-independent representation for the high dimensional data (81924 cortical thickness measures) is considerably more challenging than is the problem for any regional subset.

Moreover, the regional predictions are themselves of value, providing insight into which brain regions are related to a particular behaviour, and how strongly the measures in those regions predict that behaviour. Here we have shown that cortical thickness predicts autism symptom severity in a number of regions, and have ranked the strongest predictors. Each of these predictor regions has been associated with autism in previous research, but the much larger sample size provided by the ABIDE data lends confidence to these findings. As expected based on existing literature and given that problems with communication are part of the definition of ASD, a number of the strongest predictors are related to language: the left pars triangularis, rolandic operculum, superior temporal gyrus, and angular gyrus. The left pars triangularis is part of Broca’s area, which is critical for language production, and has been implicated in autism in numerous studies (Just et al., 2004; Zielinski et al., 2014; Lewis et al., 2014). The left rolandic operculum is involved in the production of prosody, a lack of which is one of the hallmarks of autistic speech, as well the perception of prosody, and shows abnormal levels of activiation in ASD (Paul et al., 2005; Gebauer et al., 2014). The superior temporal gyrus also does acoustic processing important for language, as well as housing Wernicke’s area, a core area for receptive language ability, and is consistently reported to show abnormalities in ASD (Lewis et al., 2014; Zilbovicius et al., 2000; Bigler et al., 2007). The angular gyrus has also been shown to be important for language (Binder et al., 1997), and to exhibit abnormalities in ASD (Just et al., 2004). Issues with social interaction is also a core feature of ASD. The superior temporal gyrus is also involved in non-language social cognition (Adolphs, 2001), as well as the adjacent superior temporal sulcus (Allison et al., 2000); both have been implicated in this domain in ASD (Di Martino et al., 2009; Zilbovicius et al., 2006; Redcay, 2008). The bilateral intraparietal sulci are also involved in social cognition. They are considered part of the mirror neuron system (Rizzolatti and Fabbri-Destro, 2010), and play a role in interpreting the intentions of the actions of others (Hamilton and Grafton, 2006). Another core aspect of social cognition is social orienting/joint attention, which has been argued to be defective in ASD (Mundy et al., 1990; Dawson et al., 2004). These aspects of social cognition have been linked to the anterior cingulate cortex and to dorsal medial frontal cortex, both of which show abnormalities in ASD (Mundy et al., 2009; Mundy, 2003). The third part of the ASD definition involves repetitive patterns of behaviour, exemplified by stereotypic body movements such as hand-flapping. Such repetitive behaviours have been suggested to relate to basal ganglia dysfunction in the inhibition of supplementary motor and motor areas (Mink, 1996).

In addition to these core behavioural abnormalities, motor and sensory processing abnormalities are pervasive in children and adults with autism (Smith, 2004; Marco et al., 2011; Leekam et al., 2007). Individuals with autism exhibit a range of motor abnormalities (Smith, 2004), and both hypo- and hyper-sensitivity to visual, auditory, and tactile inputs (Leekam et al., 2007). In this respect, it is interesting to note that some of the strongest predictors seen here are in regions associated with low level processing of motor, visual, auditory, and tactile inputs. Abnormalities in motor behaviors in ASD are associated with abnormalities in motor and supplementary motor cortex (Mostofsky et al., 2007). Visual processing involves the striate cortex within the calcarine fissure, and the surrounding cortex, including the cuneus, the caudal portion of the precuneus, and the lingual gyrus. Findings of abnormalities in visual cortex in ASD are common (Barbeau et al., 2015; Samson et al., 2012; Philip et al., 2012; Green et al., 2013). Auditory processing involves Heschl’s gyrus and the surrounding cortex within the superior temporal gyrus. Individuals with ASD have been reported to show abnormalities in these areas (O’Connor, 2012; Samson et al., 2011; Green et al., 2013). Tactile processing involves the postcentral gyrus, which also exhibits abnormalities in individuals with ASD (Rumsey et al., 1985; Horwitz et al., 1988; Kaiser et al., 2015).

A possible limitation of the study is that the severity scores that we aim to predict are integer valued with a limited range (as can be observed in Fig. 3) and therefore the continuity assumption made in the regression models might not be correct. A possible solution would be the use of the methods for ordinal regression, where the response variables are treated as ordered categories and not as continuous variables (Bender and Grouven, 1997; Chu and Keerthi, 2007). However, since the severity scores also carry metric information (Gotham et al., 2009), not used in the ordinal regression,it is unclear if ordinal regression models would be suitable for the task.

It bears repeating that the methods described here for research with multi-site, multi-protocol data are applicable to any such data. The results here served to demonstrate the validity of the methods, and their use in identifying and ranking regional brain measures as predictors of behaviour. But the brain measures need not be cortical thickness, and the predicted behavioural measures need not be the severity of symptoms of ASD.

## Footnotes

^1^http://fcon_1000.projects.nitrc.org/indi/abide/

^2^http://fcon_1000.projects.nitrc.org/indi/abide/

^3^The *Q*^2^ provides a measure of how well out-of-training set severity scores are predictable by the learned model (http://scikit-learn.org/stable/modules/model_evaluation.html#regression-metrics). It is defined as 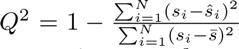, where *ŝ_i_* is the predicted severity score for subject *i*, *s_i_* is the true severity score for subject *i*, and 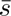 is mean of the actual/true severity scores. *Q*^2^ is bounded above by 1 but is not bounded from below. Note that *Q*^2^ does not equal *R*^2^, i.e., the correlation squared, but the *Q*^2^ value can never exceed *R*^2^. More details about different metrics and their relations are available in the supplement (Section 6).

## Acknowledgments

The authors wish to acknowledge CSC – IT Center for Science Ltd., Finland, for the allocation of computational resources. This research has been supported by The Azrieli Neurodevelopmental Research Program in partnership with Brain Canada Multi-Investigator Research Initiative (MIRI) grant to BSK, JDL and ACE. This research was enabled in part by support provided by Calcul Quebec (www.calculquebec.ca) and Compute Canada (www.computecanada.ca).

This project has received funding from the Universidad Carlos III de Madrid, the European Union’s Seventh Framework Programme for research, technological development and demonstration under grant agreement nr 600371, el Ministerio de Economía y Competitividad (COFUND2013-40258) and Banco Santander.

## References

Abdi, H., 2007. Singular value decomposition (svd) and generalized singular value decomposition. Encyclopedia of measurement and statistics. Thousand Oaks (CA): Sage, 907–912.

Adolphs, R., 2001. The neurobiology of social cognition. Current opinion in neurobiology 11, 231–239.

Allison, T., Puce, A., McCarthy, G., 2000. Social perception from visual cues: role of the sts region. Trends in cognitive sciences 4, 267–278.

Amaral, D.G., Schumann, C.M., Nordahl, C.W., 2008. Neuroanatomy of autism. Trends in neurosciences 31, 137–145.

Anderson, M.J., Robinson, J., 2001. Permutation tests for linear models. Australian & New Zealand Journal of Statistics 43, 75–88.

Auzias, G., Breuil, C., Takerkart, S., Deruelle, C., 2014. Detectability of brain structure abnormalities related to autism through mri-derived measures from multiple scanners, in: Biomedical and Health Informatics (BHI), 2014 IEEE-EMBS International Conference on, IEEE. pp. 314–317.

Auzias, G., Takerkart, S., Deruelle, C., 2016. On the influence of confounding factors in multi-site brain morphometry studies of developmental pathologies: Application to autism spectrum disorder. IEEE J Biomed Health Inform 20, 810–817.

Barbeau, E.B., Lewis, J.D., Doyon, J., Benali, H., Zeffiro, T.A., Mottron, L., 2015. A greater involvement of posterior brain areas in interhemispheric transfer in autism: fmri, dwi and behavioral evidences. NeuroImage: Clinical 8, 267–280.

Barnea-Goraly, N., Kwon, H., Menon, V., Eliez, S., Lotspeich, L., Reiss, A.L., 2004. White matter structure in autism: preliminary evidence from diffusion tensor imaging. Biological psychiatry 55, 323–326.

Bauman, M.L., 1991. Microscopic neuroanatomic abnormalities in autism. Pediatrics 87, 791–796.

Bender, R., Grouven, U., 1997. Ordinal logistic regression in medical research. Journal of the Royal College of Physicians of London 31, 546–551.

Bigler, E.D., Mortensen, S., Neeley, E.S., Ozonoff, S., Krasny, L., Johnson, M., Lu, J., Provencal, S.L., McMahon, W., Lainhart, J.E., 2007. Superior temporal gyrus, language function, and autism. Developmental neuropsychology 31, 217–238.

Binder, J.R., Frost, J.A., Hammeke, T.A., Cox, R.W., Rao, S.M., Prieto, T., 1997. Human brain language areas identified by functional magnetic resonance imaging. The Journal of Neuroscience 17, 353–362.

Boser, B.E., Guyon, I.M., Vapnik, V.N., 1992. A training algorithm for optimal margin classifiers, in: Proceedings of the fifth annual workshop on Computational learning theory, ACM. pp. 144–152.

Button, K.S., Ioannidis, J.P., Mokrysz, C., Nosek, B.A., Flint, J., Robinson, E.S., Munafo, M.R., 2013. Power failure: why small sample size undermines the reliability of neuroscience. Nature Reviews Neuroscience 14, 365–376.

Castrillon, J.G., Ahmadi, A., Navab, N., Richiardi, J., 2014. Learning with multi-site fmri graph data, in: 2014 48th Asilomar Conference on Signals, Systems and Computers, IEEE. pp. 608–612.

Cerliani, L., Mennes, M., Thomas, R.M., Di Martino, A., Thioux, M., Keysers, C., 2015. Increased functional connectivity between subcortical and cortical resting-state networks in autism spectrum disorder. JAMA psychiatry 72, 767–777.

Chang, C.C., Lin, C.J., 2011. Libsvm: A library for support vector machines. ACM Transactions on Intelligent Systems and Technology (TIST) 2, 27.

Chu, W., Keerthi, S.S., 2007. Support vector ordinal regression. Neural computation 19, 792–815.

Collins, D.L., Neelin, P., Peters, T.M., Evans, A.C., 1994. Automatic 3d intersubject registration of mr volumetric data in standardized talairach space. Journal of computer assisted tomography 18, 192–205.

Cortes, C., Vapnik, V., 1995. Support-vector networks. Machine learning 20, 273–297.

Courchesne, E., Mouton, P.R., Calhoun, M.E., Semendeferi, K., Ahrens-Barbeau, C., Hallet, M.J., Barnes, C.C., Pierce, K., 2011. Neuron number and size in prefrontal cortex of children with autism. Jama 306, 2001–2010.

Dawson, G., Toth, K., Abbott, R., Osterling, J., Munson, J., Estes, A., Liaw, J., 2004. Early social attention impairments in autism: social orienting, joint attention, and attention to distress. Developmental psychology 40, 271.

De Jong, S., 1993. Simpls: an alternative approach to partial least squares regression. Chemometrics and intelligent laboratory systems 18, 251–263.

Devlin, B., Scherer, S.W., 2012. Genetic architecture in autism spectrum disorder. Current opinion in genetics & development 22, 229–237.

Di Martino, A., Ross, K., Uddin, L.Q., Sklar, A.B., Castellanos, F.X., Milham, M.P., 2009. Functional brain correlates of social and nonsocial processes in autism spectrum disorders: an activation likelihood estimation meta-analysis. Biological psychiatry 65, 63–74.

Di Martino, A., Yan, C.G., Li, Q., Denio, E., Castellanos, F.X., Alaerts, K., Anderson, J.S., Assaf, M., Bookheimer, S.Y., Dapretto, M., et al., 2014. The autism brain imaging data exchange: towards a large-scale evaluation of the intrinsic brain architecture in autism. Molecular psychiatry 19, 659–667.

Doyle-Thomas, K.A., Duerden, E.G., Taylor, M.J., Lerch, J.P., Soorya, L.V., Wang, A.T., Fan, J., Hollander, E., Anagnostou, E., 2013. Effects of age and symptomatology on cortical thickness in autism spectrum disorders. Research in autism spectrum disorders 7, 141–150.

Ecker, C., Rocha-Rego, V., Johnston, P., Mourao-Miranda, J., Marquand, A., Daly, E.M., Brammer, M.J., Murphy, C., Murphy, D.G., Consortium, M.A., et al., 2010. Investigating the predictive value of whole-brain structural mr scans in autism: a pattern classification approach. Neuroimage 49, 44–56.

Fatemi, S.H., Halt, A.R., Realmuto, G., Earle, J., Kist, D.A., Thuras, P., Merz, A., 2002. Purkinje cell size is reduced in cerebellum of patients with autism. Cellular and molecular neurobiology 22, 171–175.

Friedman, L., Wall, M., 2005. Graphical views of suppression and multi-collinearity in multiple linear regression. The American Statistician 59, 127–136.

Gammerman, A., Vovk, V., Vapnik, V., 1998. Learning by transduction, in: Proceedings of the Fourteenth conference on Uncertainty in artificial intelligence, Morgan Kaufmann Publishers Inc.. pp. 148–155.

Gebauer, L., Skewes, J., Hørlyck, L., Vuust, P., 2014. Atypical perception of affective prosody in autism spectrum disorder. Neuroimage: Clinical 6, 370–378.

Georgiades, S., Szatmari, P., Boyle, M., Hanna, S., Duku, E., Zwaigenbaum, L., Bryson, S., Fombonne, E., Volden, J., Mirenda, P., et al., 2013. Investigating phenotypic heterogeneity in children with autism spectrum disorder: a factor mixture modeling approach. Journal of Child Psychology and Psychiatry 54, 206–215.

Gillberg, C., 1993. Autism and related behaviours. Journal of Intellectual Disability Research 37, 343–372.

Gong, B., Shi, Y., Sha, F., Grauman, K., 2012. Geodesic flow kernel for unsupervised domain adaptation, in: Computer Vision and Pattern Recognition (CVPR), 2012 IEEE Conference on, IEEE. pp. 2066–2073.

Gotham, K., Pickles, A., Lord, C., 2009. Standardizing ados scores for a measure of severity in autism spectrum disorders. Journal of autism and developmental disorders 39, 693–705.

Gotham, K., Pickles, A., Lord, C., 2012. Trajectories of autism severity in children using standardized ados scores. Pediatrics 130, e1278–e1284.

Gotham, K., Risi, S., Pickles, A., Lord, C., 2007. The autism diagnostic observation schedule: revised algorithms for improved diagnostic validity. Journal of autism and developmental disorders 37, 613–627.

Green, S.A., Rudie, J.D., Colich, N.L., Wood, J.J., Shirinyan, D., Hernandez, L., Tottenham, N., Dapretto, M., Bookheimer, S.Y., 2013. Overreactive brain responses to sensory stimuli in youth with autism spectrum disorders. Journal of the American Academy of Child & Adolescent Psychiatry 52, 1158–1172.

Gupta, C.N., Calhoun, V.D., Rachakonda, S., Chen, J., Patel, V., Liu, J., Segall, J., Franke, B., Zwiers, M.P., Arias-Vasquez, A., et al., 2015. Patterns of gray matter abnormalities in schizophrenia based on an international mega-analysis. Schizophrenia bulletin 41, 1133–1142.

Haar, S., Berman, S., Behrmann, M., Dinstein, I., 2016. Anatomical abnormalities in autism? Cerebral Cortex 26,1440–1452.

Hadjikhani, N., Joseph, R.M., Snyder, J., Tager-Flusberg, H., 2006. Anatomical differences in the mirror neuron system and social cognition network in autism. Cerebral cortex 16, 1276–1282.

Hamilton, A.F.d.C., Grafton, S.T., 2006. Goal representation in human anterior intraparietal sulcus. The Journal of Neuroscience 26, 1133–1137.

Horwitz, B., Rumsey, J.M., Grady, C.L., Rapoport, S.I., 1988. The cerebral metabolic landscape in autism: intercorrelations of regional glucose utilization. Archives of neurology 45, 749–755.

Hugdahl, K., 2005. Symmetry and asymmetry in the human brain. European Review 13, 119–133.

Jacobson, R., Le Couteur, A., Howlin, P., Rutter, M., 1988. Selective subcortical abnormalities in autism. Psychological medicine 18, 39–48.

Jiang, J., 2008. A literature survey on domain adaptation of statistical classifiers. Technical report, Computer Science Department at University of Illinois at Urbana-Champaign. Available at http://sifaka.cs.uiuc.edu/jiang4/domainadaptation/survey.

Johnson, M.H., Gliga, T., Jones, E., Charman, T., 2015. Annual research review: Infant development, autism, and adhd-early pathways to emerging disorders. Journal of Child Psychology and Psychiatry 56, 228–247.

Just, M.A., Cherkassky, V.L., Keller, T.A., Minshew, N.J., 2004. Cortical activation and synchronization during sentence comprehension in high-functioning autism: evidence of underconnectivity. Brain 127, 1811–1821.

Kaiser, M.D., Yang, D.Y.J., Voos, A.C., Bennett, R.H., Gordon, I., Pretzsch, C., Beam, D., Keifer, C., Eilbott, J., McGlone, F., et al., 2015. Brain mechanisms for processing affective (and nonaffective) touch are atypical in autism. Cerebral Cortex, bhv125.

Khundrakpam, B.S., Tohka, J., Evans, A.C., Group, B.D.C., et al., 2015. Prediction of brain maturity based on cortical thickness at different spatial resolutions. Neuroimage 111, 350–359.

Kim, J.S., Singh, V., Lee, J.K., Lerch, J., Ad-Dab’bagh, Y., MacDonald, D., Lee, J.M., Kim, S.I., Evans, A.C., 2005. Automated 3-d extraction and evaluation of the inner and outer cortical surfaces using a laplacian map and partial volume effect classification. Neuroimage 27, 210–221.

Klöppel, S., Peter, J., Ludl, A., Pilatus, A., Maier, S., Mader, I., Heimbach, B., Frings, L., Egger, K., Dukart, J., et al., 2015. Applying automated mr-based diagnostic methods to the memory clinic: A prospective study. Journal of Alzheimer’s Disease 47, 939–954.

Kostro, D., Abdulkadir, A., Durr, A., Roos, R., Leavitt, B.R., Johnson, H., Cash, D., Tabrizi, S.J., Scahill, R.I., Ronneberger, O., et al., 2014. Correction of inter-scanner and within-subject variance in structural mri based automated diagnosing. NeuroImage 98, 405–415.

Leekam, S.R., Nieto, C., Libby, S.J., Wing, L., Gould, J., 2007. Describing the sensory abnormalities of children and adults with autism. Journal of autism and developmental disorders 37, 894–910.

de Leeuw, J., 2007. Derivatives of generalized eigensystems with applications. UCLA Department of Statistics Papers, 1–28.

Lefebvre, A., Beggiato, A., Bourgeron, T., Toro, R., 2015. Neuroanatomical diversity of corpus callosum and brain volume in autism: meta-analysis, analysis of the autism brain imaging data exchange project, and simulation. Biological psychiatry 78, 126–134.

Lewis, J.D., Evans, A., Pruett, J., Botteron, K., Zwaigenbaum, L., Estes, A., Gerig, G., Collins, L., Kostopoulos, P., McKinstry, R., et al., 2014. Network inefficiencies in autism spectrum disorder at 24 months. Translational psychiatry 4, e388.

Lewis, J.D., Theilmann, R.J., Townsend, J., Evans, A.C., 2013. Network efficiency in autism spectrum disorder and its relation to brain overgrowth. Frontiers in human neuroscience 7, 845.

Lord, C., Jones, R.M., 2012. Annual research review: Re-thinking the classification of autism spectrum disorders. Journal of Child Psychology and Psychiatry 53, 490–509.

Lord, C., Risi, S., Lambrecht, L., Cook Jr, E.H., Leventhal, B.L., DiLavore, P.C., Pickles, A., Rutter, M., 2000. The autism diagnostic observation schedule—generic: A standard measure of social and communication deficits associated with the spectrum of autism. Journal of autism and developmental disorders 30, 205–223.

Lyttelton, O., Boucher, M., Robbins, S., Evans, A., 2007. An unbiased iterative group registration template for cortical surface analysis. Neuroimage 34, 1535–1544.

Marco, E.J., Hinkley, L.B., Hill, S.S., Nagarajan, S.S., 2011. Sensory processing in autism: a review of neurophysiologic findings. Pediatric Research 69, 48R–54R.

Mink, J.W., 1996. The basal ganglia: focused selection and inhibition of competing motor programs. Progress in neurobiology 50, 381–425.

Mostofsky, S.H., Burgess, M.P., Larson, J.C.G., 2007. Increased motor cortex white matter volume predicts motor impairment in autism. Brain 130, 2117–2122.

Mundy, P., 2003. Annotation: The neural basis of social impairments in autism: the role of the dorsal medial-frontal cortex and anterior cingulate system. Journal of Child Psychology and psychiatry 44, 793–809.

Mundy, P., Sigman, M., Kasari, C., 1990. A longitudinal study of joint attention and language development in autistic children. Journal of Autism and developmental Disorders 20, 115–128.

Mundy, P., Sullivan, L., Mastergeorge, A.M., 2009. A parallel and distributed-processing model of joint attention, social cognition and autism. Autism research 2, 2–21.

Nielsen, J.A., Zielinski, B.A., Fletcher, P.T., Alexander, A.L., Lange, N., Bigler, E.D., Lainhart, J.E., Anderson, J.S., 2013. Multisite functional connectivity mri classification of autism: Abide results. Frontiers in human neuroscience 7.

O’Connor, K., 2012. Auditory processing in autism spectrum disorder: a review. Neuroscience & Biobehavioral Reviews 36, 836–854.

Pan, S.J., Yang, Q., 2010. A survey on transfer learning. Knowledge and Data Engineering, IEEE Transactions on 22, 1345–1359.

Paul, R., Augustyn, A., Klin, A., Volkmar, F.R., 2005. Perception and production of prosody by speakers with autism spectrum disorders. Journal of autism and developmental disorders 35, 205–220.

Philip, R.C., Dauvermann, M.R., Whalley, H.C., Baynham, K., Lawrie, S.M., Stanfield, A.C., 2012. A systematic review and meta-analysis of the fmri investigation of autism spectrum disorders. Neuroscience & Biobehavioral Reviews 36, 901–942.

Qian, J., Hastie, T., Friedman, J., Tibshirani, R., Simon, N., 2013. Glmnet for matlab, 2013. http://www.stanford.edu/~hastie/glmnet_matlab/

Raznahan, A., Lenroot, R., Thurm, A., Gozzi, M., Hanley, A., Spence, S.J., Swedo, S.E., Giedd, J.N., 2013. Mapping cortical anatomy in preschool aged children with autism using surface-based morphometry. Neuroimage: Clinical 2, 111–119.

Raznahan, A., Lerch, J.P., Lee, N., Greenstein, D., Wallace, G.L., Stockman, M., Clasen, L., Shaw, P.W., Giedd, J.N., 2011. Patterns of coordinated anatomical change in human cortical development: a longitudinal neuroimaging study of maturational coupling. Neuron 72, 873–884.

Redcay, E., 2008. The superior temporal sulcus performs a common function for social and speech perception: implications for the emergence of autism. Neuroscience & Biobehavioral Reviews 32, 123–142.

Rizzolatti, G., Fabbri-Destro, M., 2010. Mirror neurons: from discovery to autism. Experimental Brain Research 200, 223–237.

Rojas, D.C., Peterson, E., Winterrowd, E., Reite, M.L., Rogers, S.J., Tregellas, J.R., 2006. Regional gray matter volumetric changes in autism associated with social and repetitive behavior symptoms. BMC psychiatry 6, 56.

Rosipal, R., Krämer, N., 2006. Overview and recent advances in partial least squares, in: Subspace, latent structure and feature selection. Springer, pp. 34–51.

Rumsey, J.M., Duara, R., Grady, C., Rapoport, J.L., Margolin, R.A., Rapoport, S.I., Cutler, N.R., 1985. Brain metabolism in autism: Resting cerebral glucose utilization rates as measured with positron emission tomography. Archives of general psychiatry 42, 448–455.

Samson, F., Hyde, K.L., Bertone, A., Soulières, I., Mendrek, A., Ahad, P., Mottron, L., Zeffiro, T.A., 2011. Atypical processing of auditory temporal complexity in autistics. Neuropsychologia 49, 546–555.

Samson, F., Mottron, L., Soulieres, I., Zeffiro, T.A., 2012. Enhanced visual functioning in autism: An ale meta-analysis. Human brain mapping 33, 1553–1581.

Sato, J.R., Hoexter, M.Q., de Magalhaes Oliveira, P.P., Brammer, M.J., Murphy, D., Ecker, C., Consortium, M.A., et al., 2013. Inter-regional cortical thickness correlations are associated with autistic symptoms: a machine-learning approach. Journal of psychiatric research 47, 453–459.

Schölkopf, B., Smola, A.J., Williamson, R.C., Bartlett, P.L., 2000. New support vector algorithms. Neural computation 12, 1207–1245.

Schumann, C.M., Barnes, C.C., Lord, C., Courchesne, E., 2009. Amygdala enlargement in toddlers with autism related to severity of social and communication impairments. Biological psychiatry 66, 942–949.

Shaw, P., Kabani, N.J., Lerch, J.P., Eckstrand, K., Lenroot, R., Gogtay, N., Greenstein, D., Clasen, L., Evans, A., Rapoport, J.L., et al., 2008. Neurodevelopmental trajectories of the human cerebral cortex. The Journal of Neuroscience 28, 3586–3594.

Shi, Y., Sha, F., 2012. Information-theoretical learning of discriminative clusters for unsupervised domain adaptation. In International conference on machine learning (ICML12), pages 1079–1086.

Sled, J.G., Zijdenbos, A.P., Evans, A.C., 1998. A nonparametric method for automatic correction of intensity nonuniformity in mri data. Medical Imaging, IEEE Transactions on 17, 87–97.

Smith, I.M., 2004. Motor problems in children with autistic spectrum disorders. Developmental motor disorders: A neuropsychological perspective, 152–168.

Smola, A.J., Schölkopf, B., 2004. A tutorial on support vector regression. Statistics and computing 14, 199–222.

Szatmari, P., Georgiades, S., Duku, E., Bennett, T.A., Bryson, S., Fombonne, E., Mirenda, P., Roberts, W., Smith, I.M., Vaillancourt, T., et al., 2015. Developmental trajectories of symptom severity and adaptive functioning in an inception cohort of preschool children with autism spectrum disorder. JAMA psychiatry 72, 276–283.

Tohka, J., Moradi, E., Huttunen, H., 2016. Comparison of feature selection techniques in machine learning for anatomical brain mri in dementia. Neuroinformatics 14, 279–296.

Tohka, J., Zijdenbos, A., Evans, A., 2004. Fast and robust parameter estimation for statistical partial volume models in brain mri. Neuroimage 23, 84–97.

Torgerson, C., GENDAAR Working Group, t., Irimia, A., Horn, J.V., 2015. The search for structural biomarkers in autism spectrum disorders, in: Annual Meeting of the Organisation for Human Brain Mapping.

Vapnik, V., 1995. The nature of statistical learning theory. Springer, New York.

Vapnik, V.N., Vapnik, V., 1998. Statistical learning theory. volume 1. Wiley New York.

Wang, L., Wee, C.Y., Tang, X., Yap, P.T., Shen, D., 2015. Multi-task feature selection via supervised canonical graph matching for diagnosis of autism spectrum disorder. Brain imaging and behavior, 1–8.

Webb, S.J., Jones, E.J., Merkle, K., Venema, K., Greenson, J., Murias, M., Dawson, G., 2011. Developmental change in the erp responses to familiar faces in toddlers with autism spectrum disorders versus typical development. Child development 82, 1868–1886.

Wing, L., 1997. The autistic spectrum. The lancet 350, 1761–1766.

Wolff, J.J., Gu, H., Gerig, G., Elison, J.T., Styner, M., Gouttard, S., Botteron, K.N., Dager, S.R., Dawson, G., Estes, A.M., et al., 2014. Differences in white matter fiber tract development present from 6 to 24 months in infants with autism. American Journal of Psychiatry.

Zielinski, B.A., Prigge, M.B., Nielsen, J.A., Froehlich, A.L., Abildskov, T.J., Anderson, J.S., Fletcher, P.T., Zygmunt, K.M., Travers, B.G., Lange, N., et al., 2014. Longitudinal changes in cortical thickness in autism and typical development. Brain 137, 1799–1812.

Zijdenbos, A.P., Forghani, R., Evans, A.C., 2002. Automatic” pipeline” analysis of 3-d mri data for clinical trials: application to multiple sclerosis. Medical Imaging, IEEE Transactions on 21, 1280–1291.

Zilbovicius, M., Boddaert, N., Belin, P., Poline, J.B., Remy, P., Mangin, J.F., Thivard, L., Barthélémy, C., Samson, Y., 2000. Temporal lobe dysfunction in childhood autism: a pet study. American Journal of Psychiatry 157, 1988–1993.

Zilbovicius, M., Meresse, I., Chabane, N., Brunelle, F., Samson, Y., Boddaert, N., 2006. Autism, the superior temporal sulcus and social perception. Trends in neurosciences 29, 359–366.

Zou, H., Hastie, T., 2005. Regularization and variable selection via the elastic net. Journal of the Royal Statistical Society: Series B (Statistical Methodology) 67, 301–320.

